## Supplementary material for "Human Neuronal Excitation/Inhibition Balance Explains and Predicts Neurostimulation Induced Learning Benefits": S1_Text

### Supplementary Information

#### Supplementary Introduction (osf.io/y4xar)

In addition to the behavioral aspects, overlearning recently has been suggested to be associated with a change in the ratio of excitation and inhibition (E/I ratio) [1]. The E/I ratio shifted towards a more inhibitory dominant state after overlearning, in this case in a simple visual task and superseded learning. The balance between excitation and inhibition controls the temporal organization of neuronal avalanches which can be considered as a robust feature of spontaneous neuronal activity and are approximated by a power law [2]. Human resting-state magnetoencephalography (MEG) [2] and electroencephalography (EEG) [3] consist of neuronal avalanches, suggesting that it is a critical state, which is typically measured via a branching parameter. These branching parameters indicate the degree to which a signal propagates between clusters of neurons, with a branching parameter of 1 indicating criticality. Therefore, the present study also investigated if neuronal avalanches could serve as an electrophysiological marker for arithmetic learning and overlearning, and if the effect of tRNS is predicted by the individuals' neuronal avalanches.

We reasoned that a branching parameter of neuronal avalanches (in this case, kappa, κ) different from 1 may be seen amongst overlearners, due to increased inhibition as was shown elsewhere using magnetic resonance spectroscopy [1]. Considering that tRNS increases excitation, it is expected that overlearning in combination with tRNS leads to a scaling component closer to a critical system, that is, less branching. Learning and tRNS may also impact the branching parameter due to compounded excitation following learning and active tRNS. It was also expected that learning and sham tRNS would show less branching than learning and tRNS (κ is closer to 1).

#### Supplementary Results

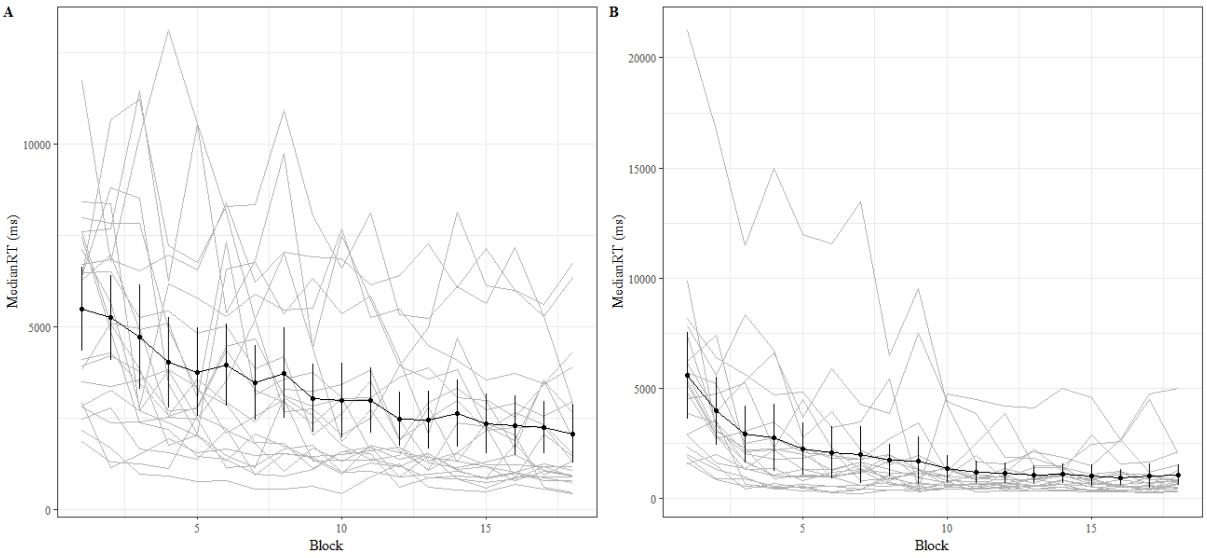


**Fig A. Individual learning curves of the RTs of the learning and overlearning task. A)** The individual learning curves of the participants who received sham stimulation (n=22) during learning shows a linear gradient. **B)** The individual learning curve of the participants who received sham stimulation (n=21) during overlearning. Bars indicate 95% confidence intervals.

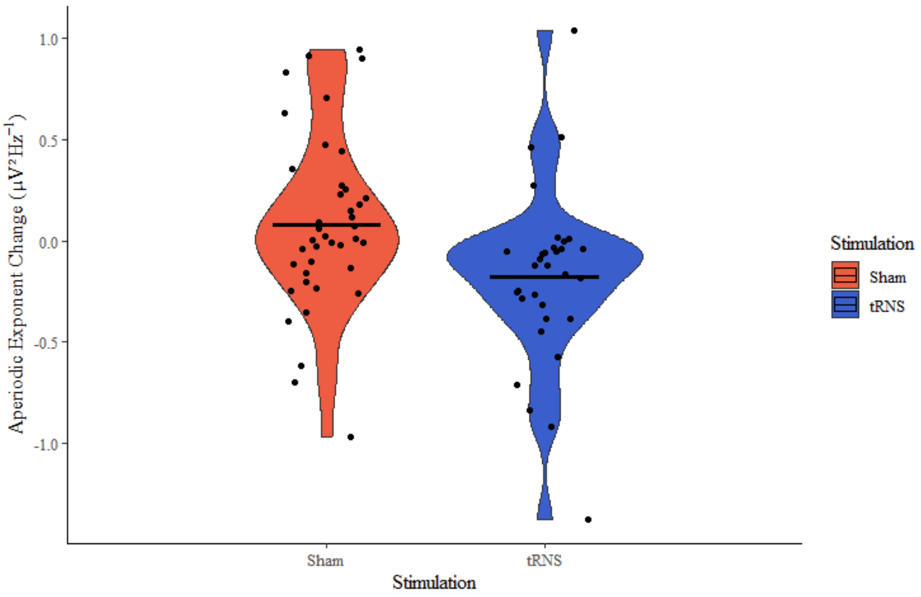


**Fig B. Individual change in aperiodic exponent for both stimulation groups.** Individual data points indicate the aperiodic exponent change (post-pre) for the sham stimulation group (in red) and the tRNS group (in blue). Means are indicated with a solid black line. Note that this figure is based after the exclusion of outliers.

##### ***Bayesian ANCOVA***

Task (learning/overlearning) and stimulation (tRNS/sham) were included as a fixed factor and the individual plateau as a covariate. The model with only stimulation included was the most probable model (P(M|data)=0.41, $BF_{M}=6.15$, $BF_{10}=3.75$) compared to the null model after observing the data. To account for model uncertainty, we looked at the Bayesian model averaging, which tested the effects of both predictors (i.e., stimulation and task) and showed that the data were 3.20 more likely under models containing stimulation as a predictor compared to all models ($BF_{incl}=3.20$). The data was only 0.35 times as likely for task as a predictor and similarly for the interaction between stimulation and task compared to all models ($BF_{incl}=0.85$). This complementary analysis strengthens the conclusions that tRNS impacts the aperiodic exponent (mean change effect=-0.11, 95% credible interval (CrI; posterior distribution that contains 95% of the data)) tRNS [-0.21, -0.01], 95% CrI Sham [0.01, 0.21]), while task has no effect (mean change effect=0.02, 95% CrI learning [-0.12, 0.06], 95% CrI overlearning [-0.07, 0.12]).

##### ***Model Comparison for Learning and Overlearning (Accuracy)***

Similarly to our brms models with RTs, we implemented several model comparisons with the leave-one-out cross-validation (LOO) to determine the best fit for the learning and overlearning task data. Note that this additional analysis served as a check since we do not expect any reliable results due to the instructions that were given to the participants. Namely, the participants were urged to avoid errors and there was no time limit present. This means that the accuracy should be consistent during the task.

Syntax:

**Mod1.A <-** brm(RTACC ~ Trial * Task + (1 + Trial|p|Participant), family=bernoulli(), iter=4000, warmup=2000)

Next, we added levels of complexity by adding our predictors stimulation and baseline aperiodic exponent to see if the fit with our data increases.

**Mod2.A <-** brm(RTACC ~ Stimulation + Trial * Task + Aperiodic_Baseline + (1 + Trial|p|Participant), family= bernoulli(), iter=4000, warmup=2000)

**Mod3.A <-** brm(RTACC ~ Trial + Task * Stimulation * Aperiodic_Baseline + (1 + Trial|p|Participant), family= bernoulli(), iter=4000, warmup=2000)

**Mod4.A <-** brm(RTACC ~ Trial * Task * Stimulation * Aperiodic_Baseline + (1 + Trial|p|Participant), family= bernoulli(), iter=4000, warmup=2000)

**Mod5.A <-** brm(RTACC ~ Trial + Task + Stimulation + Aperiodic_Baseline + (1 + Trial|p|Participant), family= bernoulli(), iter=4000, warmup=2000)

**Mod6.A <-** brm(RTACC ~ Trial * Aperiodic_Baseline * Stimulation + Task + (1 + Trial |p|Participant), family= bernoulli(), iter=4000, warmup=2000)

**Mod7.A <-** brm(RTACC ~ Stimulation * Trial + Aperiodic_Baseline + Task + (1 + Trial |p|Participant, family= bernoulli(), iter=4000, warmup=2000)

**Mod8.A <-** brm(RTACC ~ Trial + Task + Stimulation * Aperiodic_Baseline + (1 + Trial|p|Participant), family= bernoulli(), iter=4000, warmup=2000)

**Mod9.A <-** brm(RTACC ~ Trial * Aperiodic_Baseline + Task + Stimulation + (1 + Trial |p|Participant), family= bernoulli(), iter=4000, warmup=2000)

**Mod10.A <-** brm(RTACC ~ Trial + Task * Stimulation + Aperiodic_Baseline + (1 + Trial |p|Participant), family= bernoulli(), iter=4000, warmup=2000)

**Mod11.A <-** brm(RTACC ~ Trial * Task * Stimulation + Aperiodic_Baseline + (1 + Trial |p|Participant), family= bernoulli(), iter=4000, warmup=2000)

Loo(Mod1.A, Mod2.A, Mod3.A, Mod4.A, Mod5.A, Mod6.A, Mod7.A, Mod8.A, Mod9.A, Mod10.A, Mod11.A)

**Table A|** Outcome of the model comparisons in terms of predictive value for accuracy

| Model | elpd_diff | se_diff |
|---|---|---|
| Mod8.A | 0.0 | 0.0 |
| Mod5.A | -0.4 | 0.2 |
| Mod9.A | -0.7 | 0.5 |
| Mod7.A | -0.7 | 0.5 |
| Mod3.A | -0.8 | 0.7 |
| Mod10.A | -0.8 | 0.3 |
| Mod1.A | -0.9 | 0.9 |
| Mod2.A | -1.1 | 0.6 |
| Mod6.A | -1.3 | 0.6 |
| Mod11.A | -2.0 | 1.5 |
| Mod4.A | -5.0 | 1.9 |

***Model Comparisons for Learning and Overlearning (RTs)***

We implemented several model comparisons with the leave-one-out cross-validation (LOO) based on our brms model to determine the best fit for the learning and overlearning task data. First, we determined our starting model with the thought that the RTs should decrease over trials and be dependent on the task paradigm. ‘

Syntax for the priors:

prior <- c(prior("normal(0,2.5)", class = "b"), prior("normal(0,2.5)", dpar = "ndt", class = "Intercept"))

Syntax:

**Mod1** <- brm(bf(RT ~ Trial * Task + (1 + Trial|p|Participant), ndt ~ Task + (1|p|Participant), family=shifted_lognormal(), iter=4000, warmup=2000)

Next, we added levels of complexity by adding our predictors stimulation and baseline aperiodic exponent to see if the fit with our data increases.

**Mod2 <-** brm(bf(RT ~ Stimulation + Trial * Task + Aperiodic_Baseline + (1 + Trial|p|Participant), ndt ~ Task +(1|p|)) family= shifted_lognormal(), iter=4000, warmup=2000)

**Mod3 <-** brm(bf(RT ~ Trial + Task * Stimulation * Aperiodic_Baseline + (1 + Trial|p|Participant), ndt ~ Task +(1|p|Participant)), family=shifted_lognormal(), iter=4000, warmup=2000)

**Mod4 <-** brm(bf(RT ~ Trial * Task * Stimulation * Aperiodic_Baseline + (1 + Trial|p|Participant), ndt ~ Task +(1|p|Participant)), family=shifted_lognormal(), iter=4000, warmup=2000)

**Mod5 <-** brm(bf(RT ~ Trial + Task + Stimulation + Aperiodic_Baseline + (1 + Trial|p|Participant), ndt ~ Task +(1|p|Participant)), family= shifted_lognormal(), iter=4000, warmup=2000)

**Mod6 <-** brm(bf(RT ~ Trial * Aperiodic_Baseline * Stimulation + Task + (1 + Trial |p|Participant), ndt ~ Task +(1|p|Participant)), family= shifted_lognormal(), iter=4000, warmup=2000)

**Mod7 <-** brm(bf(RT ~ Stimulation * Trial + Aperiodic_Baseline + Task + (1 + Trial |p|Participant), ndt ~ Task +(1|p|Participant)), family= shifted_lognormal(), iter=4000, warmup=2000)

**Mod8 <-** brm(bf(RT ~ Trial + Task + Stimulation * Aperiodic_Baseline + (1 + Trial|p|Participant), ndt ~ Task +(1|p|Participant)), family= shifted_lognormal(), iter=4000, warmup=2000)

**Mod9 <-** brm(bf(RT ~ Trial * Aperiodic_Baseline + Task + Stimulation + (1 + Trial |p|Participant), ndt ~ Task +(1|p|Participant)), family= shifted_lognormal(), iter=4000, warmup=2000)

**Mod10 <-** brm(bf(RT ~ Trial + Task * Stimulation + Aperiodic_Baseline + (1 + Trial |p|Participant), ndt ~ Task +(1|p|Participant)), family= shifted_lognormal(), iter=4000, warmup=2000)

**Mod11 <-** brm(bf(RT ~ Trial * Task * Stimulation + Aperiodic_Baseline + (1 + Trial |p|Participant), ndt ~ Task +(1|p|Participant)), family= shifted_lognormal(), iter=4000, warmup=2000)

Loo(Mod1, Mod2, Mod3, Mod4, Mod5, Mod6, Mod7, Mod8, Mod9, Mod10, Mod11)

**Table B|** Outcome of the model comparisons in terms of predictive value for RTs

| Model | elpd_diff | ee_diff |
|---|---|---|
| Mod3 | 0.0 | 0.0 |
| Mod1 | -1.0 | 1.2 |
| Mod5 | -1.2 | 0.8 |
| Mod10 | -1.5 | 0.8 |
| Mod2 | -1.5 | 1.2 |
| Mod8 | -1.5 | 0.7 |
| Mod9 | -1.9 | 1.0 |
| Mod6 | -1.9 | 0.9 |
| Mod11 | -2.1 | 1.5 |
| Mod7 | -2.4 | 1.0 |
| Mod4 | -3.3 | 1.5 |

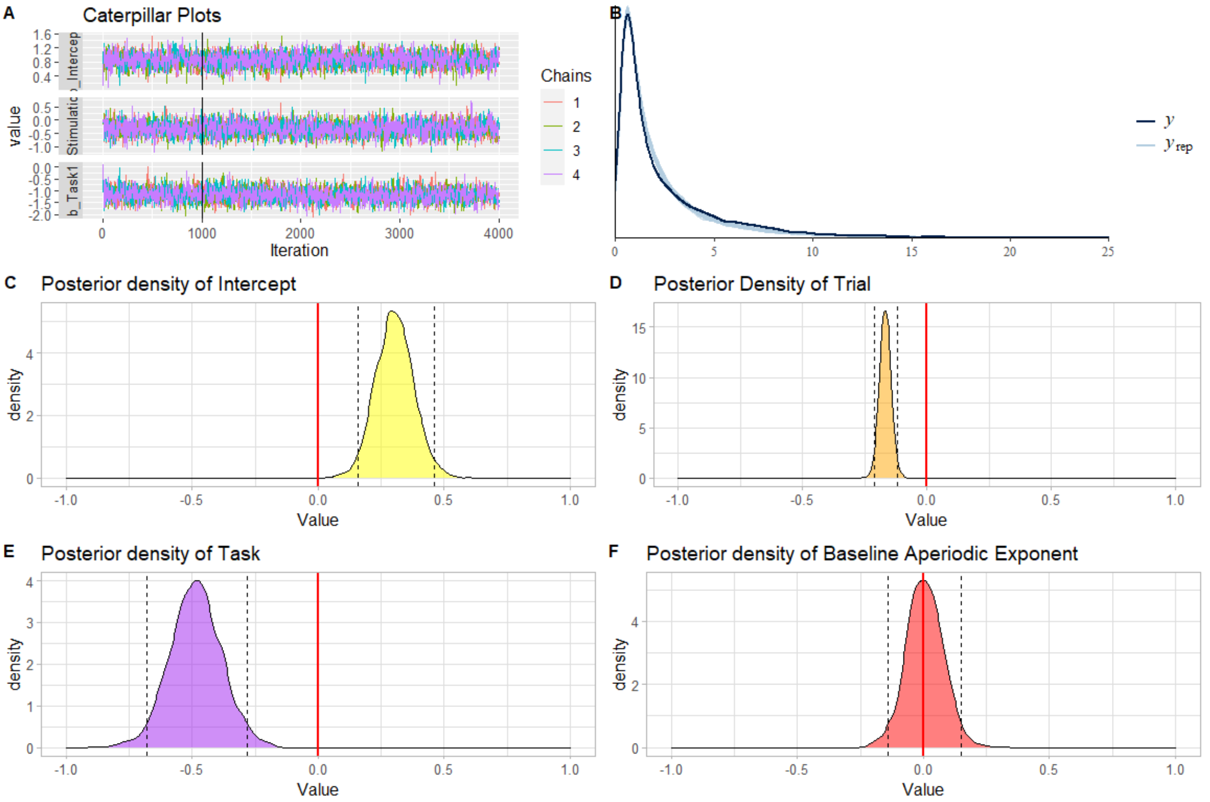


**Fig C. Output of the Bayesian mixed effects model with the four-way interaction. A)** The hairy caterpillar plots showing that convergence was reached in all four chains. **B)** Comparison of the observed outcomes (y) and the kernel density estimate of the replications of y from the posterior predictive distribution ($y_{rep}$). This posterior predictive check shows a good fit. **C)** The posterior density of the intercept. **D)** The posterior density of trial. **E)** The posterior density of task. **F)** The posterior density of the baseline aperiodic exponent.

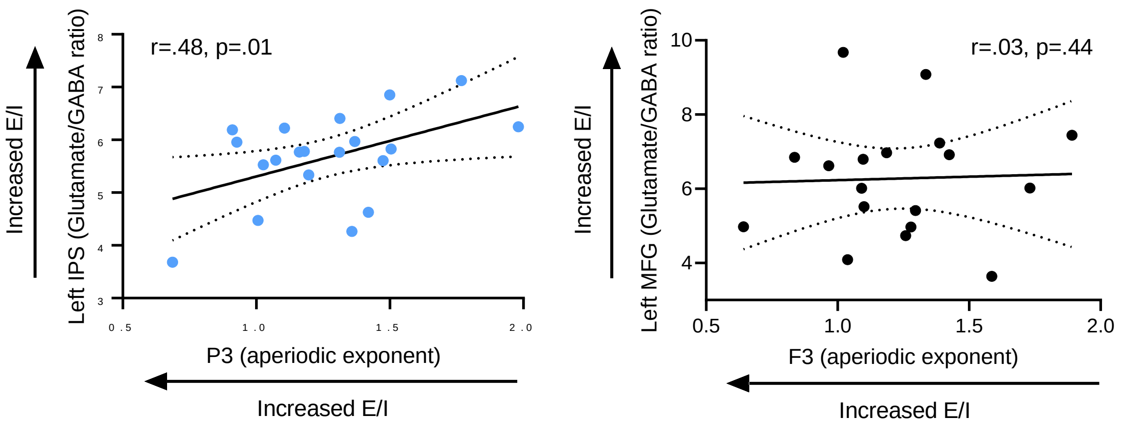


**Fig D. Correlations between MRS-based E/I and EEG-based E/I.** On the left a positive correlation between the left IPS and electrode P3, which is placed approximately above this region, showing that an increase in E/I in the IPS as it is based on glutamate/GABA is associated with a decreased E/I as indicated by aperiodic exponent ($r$=0.48, 95% CI [0.05, 0.76], $p$=.01 (one-tailed)). On the right, a non-significant correlation between the left MFG E/I and the electrode nearly this region ($r$=0.03, 95% CI [-0.44, 0.49], $p$=.44 (one-tailed)). These results are in line with our prediction that both measures characterize different aspects of E/I, and in contrast to the view that both measures reflect a similar quantification of E/I, which should have been characterized by a negative correlation.

***Neuronal Avalanches (osf.io/y4xar)***

When comparing the presence of neuronal avalanches for the different groups for the pre and the post resting-state (rs) EEG, the neuronal avalanches were plotted against a standard power law (see **Fig E**). There is no difference between the four groups regarding the pre rs-EEG. However, for the post rs-EEG there is a small diversion in the power law for the overlearning X tRNS condition. This diversion was explored further in the statistical regression analysis using κ for post rs-EEG values during overlearning. Note that we removed an additional 4 outliers in the overlearning group compared to the sample in the main manuscript. Predictors included median RT baseline, stimulation (tRNS x sham), plateau (i.e. amount of overlearning), and branching (κ) values in the pre rs-EEG and learning rate. We found that κ values during pre rs-EEG significantly predicted the κ values during post rs-EEG ($\beta$=.05$^{-1}$, *SE*=0.02$^{-1}$, *t*(27)=2.17, *p*=0.03). The predictor model was able to account for 10% of the variance of κ values in the post rs-EEG (*F*(5,27)=1.72, *p*=.16, $R^2$= .10). However, no interaction with stimulation and individual plateau was found ($\beta$=.05$^{-1}$, *SE*=0.05$^{-1}$, *t*(27)=.98 , *p*=.33).

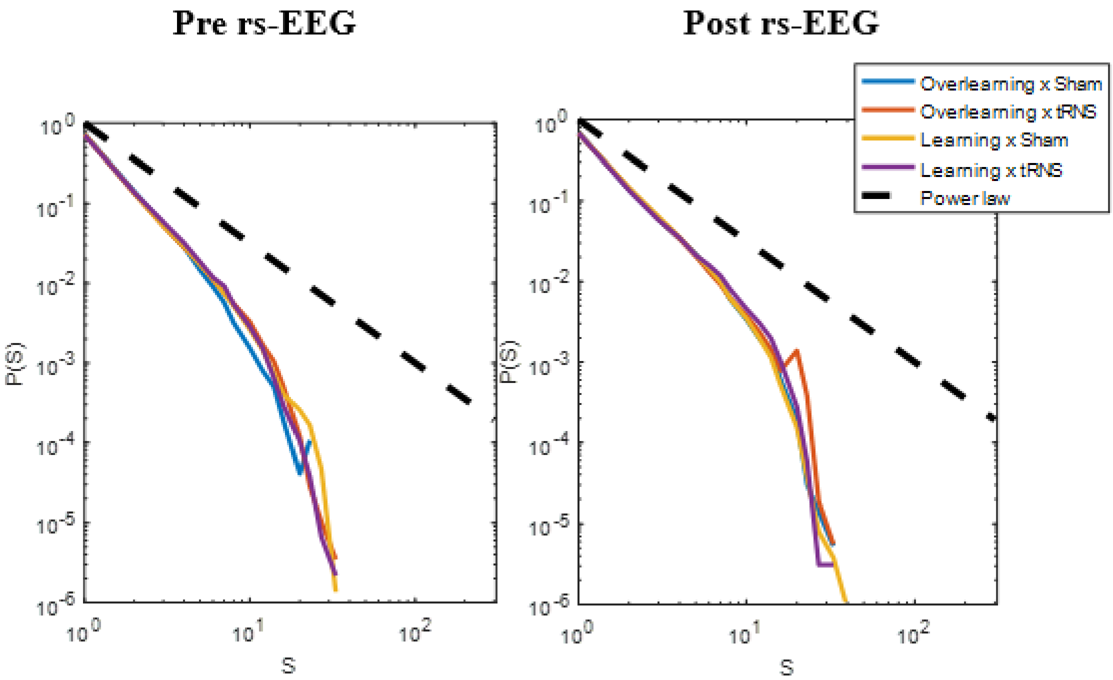


**Fig E. Neuronal avalanches for the pre and post rs-EEG for the four conditions following power laws.** Cascade size distributions are shown on the x-axis plotted against the probability on the y-axis for the pre rs-EEG (left) and the post rs-EEG (right) using $\underline{\Delta}$t=6 ms. The dashed black line represents a perfect power law with an exponent of -3/2. The different line colors in both plots indicate the four condition (task: learning x overlearning; stimulation: tRNS x sham).

**Table C|** Sensations between the tRNS and the sham stimulation group as tested with the Mann-Whitney's U test (n=102)

| Sensations | U-value | p |
|---|---|---|
| Itching | 1058 | .43 |
| Pain | 1142.5 | .92 |
| Burning | 1090.5 | .41 |
| Warmth/Heat | 1109.5 | .70 |
| Pinching | 1089 | .52 |
| Iron Taste | 1125 | .30 |
| Fatigue | 1092.5 | .74 |
| Subjective performance | 1080 | .48 |

***Model fit Quality of the Aperiodic Exponent***

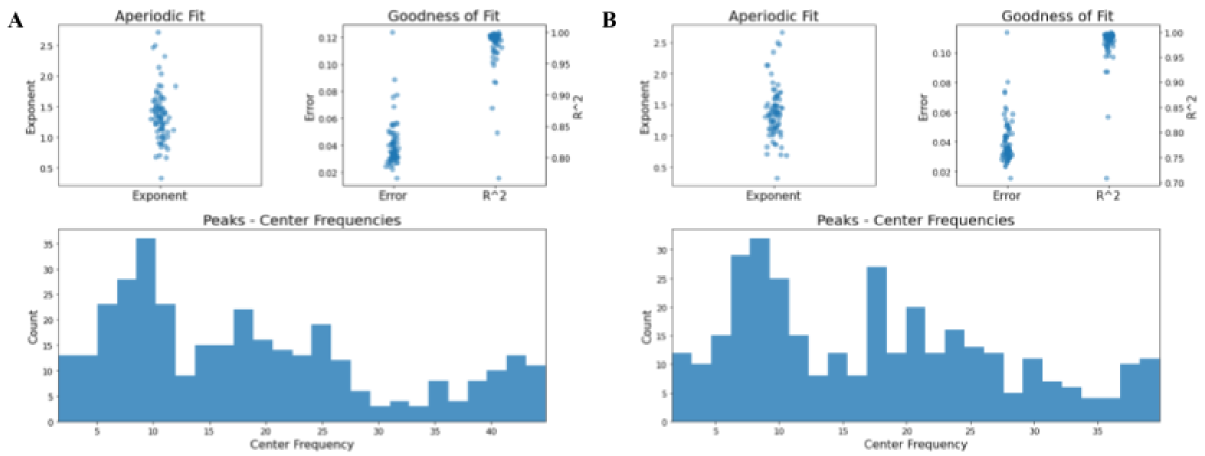


**Fig F. Raw exponent values for the baseline and post measurement. A)** Individual baseline exponent values are indicated in the top left panel. The top right panel indicates the error of the fit and the $R^2$. This plot shows a small error (mean=.04) and a high goodness of fit (mean $R^2$=.97). **B)** Individual post measurement exponent values are indicated in the top left panel. The top right panel indicates the error of the fit and the $R^2$. This plot shows a small error (mean=.11) and a high goodness of fit (mean $R^2$=.96).

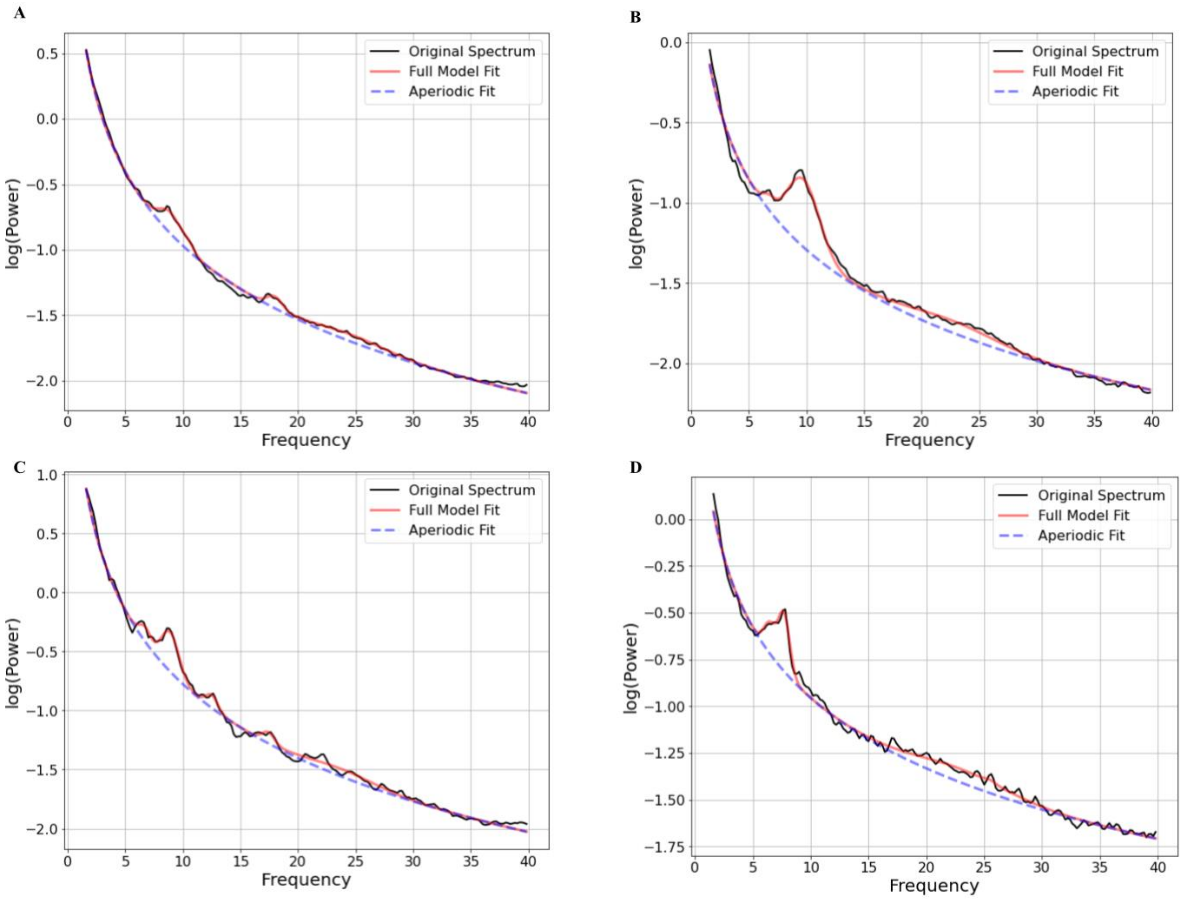


**Fig G. Averaged power spectra plotted for the separate conditions at baseline. A)** Individual power spectra averaged over the participants in the learning task who received sham stimulation (n=22). **B)** Individual power spectra averaged over the participants in the learning task who received tRNS (n=16). **C)** Individual power spectra averaged over the participants in the overlearning task who received sham stimulation (n=21). **D)** Individual power spectra averaged over the participants in the overlearning task who received tRNS (n=16).

***Transfer and Recall Performance Related to the Aperiodic Exponent (osf.io/y4xar)***

To assess any possible effects of the baseline aperiodic exponent (i.e., E/I levels) on the performance in the transfer and recall task, we ran several regression models. First, we predicted the change in median RTs between the first block of the (over)learning task and the transfer task by using a subtraction (**Table D**). We also subtracted the first block of the (over)learning task from the transfer task for the proportion of retrieval (**Table E**). Thirdly, we calculated the change in median RTs between the last block of the (over)learning task and the recall task (**Table F**), and similarly for the proportion of retrieval in the recall task (**Table G**). Predictors included the stimulation (tRNS/sham), baseline aperiodic exponent, and we corrected for the task (e.g., learning or overlearning). Additionally, we ran the same models but replaced the baseline aperiodic exponent with the change in the exponent from the pre to the post rs-EEG (**Tables H**, **I**, **J**, and **K**). No significant effects were found related to the aperiodic exponent, indicating no evidence for the importance of E/I in the transfer and recall of mathematical skills.

**Table D|** Regression for predicting the change in performance between the first block and the transfer task (median RTs) including the baseline aperiodic exponent as predictor

| **Predictors** | **Estimates** | **SE** | **CI (95%)** | **t-value** | **p-value** |
|---|---|---|---|---|---|
| (Intercept) | -.05 | .20 | -0.46 – 0.36 | -0.24 | .80 |
| Stimulation | .29 | .24 | -0.20 – 0.78 | 1.16 | .24 |
| Baseline Exponent | .08 | .19 | -0.30 – 0.47 | 0.43 | .66 |
| Task | -.11 | .24 | -0.60 – 0.37 | -0.46 | .64 |
| Stimulation X Baseline Exponent | -.21 | .25 | -0.71 – 0.30 | -0.80 | .42 |

*Note.* ***p** < 0.05; ****p** <0.01.

**Table E|** Regression for predicting the change in performance between the first block and the transfer task (proportion of retrieval) including the baseline aperiodic exponent as predictor

| Predictors | Estimates | SE | CI (95%) | t-value | p-value |
|---|---|---|---|---|---|
| (Intercept) | -.72 | .25 | -1.23 – -0.22 | -2.87 | .005** |
| Stimulation | .92 | .29 | 0.33 – 1.51 | 3.12 | .003** |
| Baseline Exponent | -.31 | .24 | -0.79 – 0.18 | -1.25 | .21 |
| Task | .49 | .29 | -0.09 – 1.08 | 1.69 | .09 |
| Stimulation X Baseline Exponent | -.03 | .32 | -0.67 – 0.61 | -0.09 | .92 |

*Note.* *$p < 0.05$; **$p < 0.01$.

**Table F|** Regression for predicting the change in performance between the last block and the recall task (median RTs) including the baseline aperiodic exponent as predictor

| Predictors | Estimates | SE | CI (95%) | t-value | p-value |
|---|---|---|---|---|---|
| (Intercept) | -.12 | .12 | -0.36 – 0.12 | -1.00 | .32 |
| Stimulation | .26 | .14 | -0.02 – 0.55 | 1.82 | .07 |
| Baseline Exponent | -.04 | .11 | -0.26 – 0.19 | -0.31 | .75 |
| Task | -.01 | .14 | -0.30 – 0.27 | -0.10 | .91 |
| Stimulation X Baseline Exponent | .12 | .14 | -0.18 – 0.41 | 0.78 | .43 |

*Note.* *$p < 0.05$; **$p < 0.01$.

**Table G|** Regression for predicting the change in performance between the last block and the recall task (proportion of retrieval) including the baseline aperiodic exponent as predictor

| Predictors | Estimates | SE | CI (95%) | t-value | p-value |
|---|---|---|---|---|---|
| (Intercept) | -.42 | .29 | -1.00 – 0.16 | -1.43 | .15 |
| Stimulation | .16 | .34 | -0.52 – 0.84 | 0.47 | .63 |
| Baseline Exponent | -0.29 | .28 | -0.85 – 0.28 | -1.01 | .31 |
| Task | .42 | .33 | -0.26 – 1.10 | 1.24 | .21 |
| Stimulation X Baseline Exponent | .27 | .37 | -0.47 – 1.01 | 0.72 | .47 |

*Note.* *$p < 0.05$; **$p < 0.01$.

**Table H|** Regression for predicting the change in performance between the first block and the transfer task (median RTs) including the aperiodic change as predictor

| Predictors | Estimates | SE | CI (95%) | t-value | p-value |
|---|---|---|---|---|---|
| (Intercept) | -.12 | .20 | -0.54 – 0.30 | -0.57 | .56 |
| Stimulation | .30 | .25 | -0.20 – 0.81 | 1.20 | .23 |
| Baseline Exponent | .10 | .17 | -0.24 – 0.45 | 0.60 | .54 |
| Task | -.06 | .24 | -0.55 – 0.43 | -0.23 | .81 |
| Stimulation X Baseline Exponent | -.10 | .25 | -0.61 – 0.42 | -0.36 | .71 |

*Note.* $^{*}p < 0.05$; $^{**}p < 0.01$.

**Table I|** Regression for predicting the change in performance between the first block and the transfer task (proportion of retrieval) including the aperiodic change as predictor

| Predictors | Estimates | SE | CI (95%) | t-value | p-value |
|---|---|---|---|---|---|
| (Intercept) | -.78 | .25 | -1.29 – -0.28 | -3.12 | .003** |
| Stimulation | .90 | .30 | 0.30 – 1.51 | 2.97 | .004** |
| Baseline Exponent | .36 | .20 | -0.05 – 0.77 | 1.75 | .08 |
| Task | .57 | .29 | -0.01 – 1.17 | 1.95 | .05 |
| Stimulation X Baseline Exponent | -.32 | .31 | -0.95 – 0.30 | -1.03 | .30 |

*Note.* $^{*}p < 0.05$; $^{**}p < 0.01$.

**Table J|** Regression for predicting the change in performance between the last block and the recall task (median RTs) including the aperiodic change as predictor

| Predictors | Estimates | SE | CI (95%) | t-value | p-value |
|---|---|---|---|---|---|
| (Intercept) | -.14 | .12 | -0.38 – 0.10 | -1.14 | .25 |
| Stimulation | .26 | .14 | -0.04 – 0.55 | 1.75 | .08 |
| Baseline Exponent | .07 | .09 | -0.12 – 0.27 | 0.73 | .46 |
| Task | $-.02e^{-1}$ | .14 | -0.29 – 0.28 | -0.01 | .98 |
| Stimulation X Baseline Exponent | -.19 | .14 | -0.48 – 0.11 | -1.23 | .22 |

*Note.* $^{*}p < 0.05$; $^{**}p < 0.01$.

**Table K|** Regression for predicting the change in performance between the last block and the recall task (proportion of retrieval) including the aperiodic change as predictor

| **Predictors** | **Estimates** | **SE** | **CI (95%)** | **t-value** | **p-value** |
|---|---|---|---|---|---|
| (Intercept) | -.41 | .29 | -0.99 – 0.16 | -1.42 | .15 |
| Stimulation | .13 | .34 | -0.57 – 0.83 | 0.36 | .71 |
| Baseline Exponent | .22 | .23 | -0.26 – 0.69 | 0.91 | .36 |
| Task | .43 | .34 | -0.24 – 1.11 | 1.27 | .20 |
| Stimulation X Baseline Exponent | -.29 | .35 | -1.00 – 0.43 | -0.79 | .43 |

*Note*. *$p < 0.05$; **$p < 0.01$.

#### Supplementary Materials and Methods

**Table L|** List of the 4 presented multiplication problems used as training and the 10 multiplication problems presented during the baseline task

| Operand 1 | Operand 2 |
|---|---|
| 15 | 5 |
| 18 | 5 |
| 12 | 5 |
| 17 | 5 |
| 23 | 4 |
| 19 | 4 |
| 14 | 3 |
| 13 | 3 |
| 27 | 2 |
| 13 | 7 |
| 26 | 2 |
| 16 | 3 |
| 18 | 4 |
| 21 | 3 |

**Table M|** List of one block of the presented multiplication problems used during the learning task and overlearning task

| Learning task | | Overlearning task | |
|---|---|---|---|
| Operand 1 | Operand 2 | Operand 1 | Operand 2 |
| 17 | 4 | 17 | 4 |
| 14 | 6 | 14 | 6 |
| 29 | 2 | 29 | 2 |
| 17 | 3 | 17 | 3 |
| 24 | 3 | 24 | 3 |
| 12 | 8 | | |
| 29 | 3 | | |
| 13 | 6 | | |
| 16 | 6 | | |
| 12 | 7 | | |

##### *Neuronal Avalanches Computation and Statistical Analysis (osf.io/y4xar)*

Due to the comparison of neural activity between the pre and post rs-EEG, we decided to remove an additional three participants from the neuronal avalanches analysis with an excluded post rs-EEG recording according to our previous exclusion criteria. To identify neuronal avalanches, standardized z-scores were calculated for each channel. Hereafter, timepoints were identified in which each channel exceeded a z score of three standard deviations (our predefined threshold) [2]. In other words, periods were identified in which each electrode contained elevated activity. Data was subsequently binned into time blocks of 6 ms. A neuronal avalanche is considered to be any length of time in which a superthreshold event occurs. In short, when 6 ms passes and no further events occur, the neuronal avalanche is over. The size of the neuronal avalanches is the number of superthreshold events (spikes) occurring before the 6 ms of time when no events is seen. Lastly, the size of the neuronal avalanches was computed, i.e., occurrence of the amount of spikes in the signal after a 6 ms time window [2,4]. Regarding the statistical analysis of neural avalanches, the post rs-EEG was compared to the pre rs-EEG (baseline) and compared between the four conditions by means of a graphical illustration and a regression model regarding $\kappa$. Branching in neuronal avalanches was indexed using $\kappa$. The dependent variable included the branching ($\kappa$) for the post rs-EEG. Predictors included the baseline performance, $\kappa$ for the pre rs-EEG, stimulation, and learning rate.

##### *Material and Methods MRS*

We recruited 22 healthy participants (16 males, mean age=26.05, standard deviation =6.5) who completed an MRI scan and EEG session for two different studies. All participants provided written, informed consent and the study was approved by the University of Oxford’s Medical Sciences Interdivisional Research Ethics Committee (MS-IDREC-C2_2015_016).

###### *MR data Acquisition and Pre-processing*

All MRI data were acquired at the Oxford Centre for Functional MRI of the Brain (FMRIB) on a 3T Siemens MAGNETOM Prisma MRI System equipped with a 32 channel receive-only head coil. Anatomical high-resolution T1-weighted scans were first acquired (MPRAGE sequence: TR=1900ms; TE=3.97ms; 192 slices; voxel size=1×1×1mm).

For MRS, spectra were measured with a semi-adiabatic localization by adiabatic selective refocusing (semi-LASER) sequence (TE=32 ms; TR=3.5 s; 32 averages) [5,6] with variable power RF pulses with optimized relaxation delays (VAPOR), water suppression and outer volume saturation. Unsuppressed water spectra acquired from the same volume of interest were used to remove residual eddy current effects and to reconstruct the phased array spectra with MRspa (https://www.cmrr.umn.edu/downloads/mrspa/). Two 20mm$^3$ voxels of interest were manually placed centred on the left intraparietal sulcus (IPS) and centred on the left inferior/middle frontal gyrus (FG) based on the individual's T1-weighted image while the participant lay down in the MR scanner. Acquisition time per voxel of interest was 10-15 minutes including sequence planning and shimming and B0 shimming.

Neurochemicals were quantified with an LCmodel [7] using a basis set of simulated spectra generated based on previously reported chemical shifts and coupling constants based on a VeSPA (versatile simulation, pulses, and analysis) simulation library [8]. Simulations were performed using the same RF pulses and sequence timings as in the 3T system described above. Absolute neurochemical concentrations were extracted from the spectra using a water signal as an internal reference.

As in previous studies, the exclusion criteria for data was the Cramér-Rao bounds [9]. Neurotransmitters quantified with Cramér-Rao lower bounds (CRLB, the estimated error of the neurotransmitter quantification) >50% were classified as not detected. Additionally, we excluded cases with an SNR beyond 3 standard deviations (per voxel of interest, per neurotransmitter), and neurotransmitter or WM capacity score that fallen beyond 3 standard deviations from the group mean. This led to the exclusion of 2 cases for the GABA measure of the frontal gyrus. For each participant, we calculated 4 (brain region (frontal, parietal) * neurochemical (GABA, glutamate)) neurotransmitter concentrations all of which were calculated as the ratios between the absolute neurotransmitter concentrations divided by the absolute concentration of total creatine (creatine+phosphocreatine). The neurotransmitter concentrations were referenced to total creatine for (i) creatine is a commonly used as a reference and it is widely accepted as an internal reference standard, (ii) its signal shares the same imperfections (e.g., frequency drift, phase drift, and subject motion) as the signal of the GABA and glutamate as all concentrations are acquired simultaneously [10] measure was similar to our description in the main text. For the correlation analysis one datapoint was defined as an outlier (±3SD from the mean) and was removed from the analysis.

GABA and glutamate levels to cognitive skill acquisition during development. *Human Brain Mapping*, *36*(11), 4334–4345. https://doi.org/10.1002/HBM.22921
